# RedOak: a reference-free and alignment-free structure for indexing a collection of similar genomes

**DOI:** 10.1101/2020.12.19.423583

**Authors:** Clément Agret, Annie Chateau, Gaetan Droc, Gautier Sarah, Manuel Ruiz, Alban Mancheron

**Affiliations:** LIRMM, Univ Montpellier, CNRS, Montpellier, France; Cirad, UMR AGAP, Avenue Agropolis, Montpellier, France; INRA, UMR AGAP, 2 Place Pierre Viala, Montpellier, France; Institut de Biologie Computationnelle, Montpellier, France; CRIStAL, Centre de Recherche en Informatique Signal et Automatique de Lille, Lille, France

## Abstract

**Background:** As the cost of DNA sequencing decreases, high-throughput sequencing technologies become increasingly accessible to many laboratories. Consequently, new issues emerge that require new algorithms, including tools for indexing and compressing hundred to thousands of complete genomes.

**Results:** This paper presents RedOak, a reference-free and alignment-free software package that allows for the indexing of a large collection of similar genomes. RedOak can also be applied to reads from unassembled genomes, and it provides a nucleotide sequence query function. This software is based on a *k*-mer approach and has been developed to be heavily parallelized and distributed on several nodes of a cluster. The source code of our RedOak algorithm is available at https://gite.lirmm.fr/doccy/RedOak.

**Conclusions:** RedOak may be really useful for biologists and bioinformaticians expecting to extract information from large sequence datasets.

## 1 Background

### Context

Complete genomes, or at least a set of sequences representing whole genomes, *i.e*.., draft genomes, are becoming increasingly easy to obtain through the intensive use of high-throughput sequencing. A new genomic era is coming, therein not only being focused on the analyses of specific genes and sequences regulating them but moving toward studies using from ten to several thousands of complete genomes per species. Such a collection is usually called a pan-genome [1, 2]. Within pan-genomes, large portions of genomes are shared between individuals. This feature could be exploited to reduce the storage cost of the genomes.

Based on this idea, this paper introduces an efficient data structure to index a collection of similar genomes in a reference- and alignment-free manner. A reference-free and alignment-free approach avoids the loss of information about genetic variation not found in the direct mapping of short sequence reads onto a reference genome [1]. Furthermore, the method presented in this paper can be applied to next-generation sequencing (NGS) reads of unassembled genomes. The method enables the easy and fast exploration of the presence-absence variation (PAV) of genes among individuals without needing the time-consuming step of *de novo* genome assembly nor the step of mapping to a reference sequence.

### Related work

One of the most commonly used data structures for genome indexing is the FM-index [3]. This compressed structure exploits the Burrows-Wheeler Transform (BWT) data reorganization properties [4] and its link with the suffix array data structure (SA) [5], which enables the construction of a genome index in linear time and space according to the genome size. To index a collection of similar genomes, J. Sirén [6] proposed creating as many BWT indexes as genomes and merging them. However, in this approach, updating the whole index seems to be a crippling obstacle because it requires merging again.

Deorowicz *et al*. [7] proposed an efficient method to store large collections of genomes. Their method uses a reference sequence and a table containing the variations of each genome from the reference, assuming that many variations are shared across the set of genomes. The data structure is then compressed, enabling the efficient storage of a set of very close genomic sequences. Their structure cannot be queried, and retrieving a genome consists of decompressing the data and applying the indexed variations to the reference sequence.

New methods have emerged for both the storage and analysis of pan-genomes. These methods usually use the same approach, which consists of storing a reference sequence and information on each genome variation compared to the reference. This implies that such tools require prior information on genomic variations. This information must have been previously computed, for example, by performing a multiple-genome alignment. Some of these methods, such as SplitMEM [8] and TwoPaCo [9], use graphs or combine graphs with a generalized compressed suffix array (GCSA) [10, 11, 12]. Other methods use a custom data structure based on sequence alignment methods [13, 14]. MuGI [15] stores the reference in compact form (4 bits to encode a single char), a variant database (one bit vector for each variant), and an array retaining information about each *k*-mer.

Few methods aim to store a pan-genome without prior knowledge, and even fewer methods allow direct query on pan-genomes. CHICO [16] program uses a hybrid index that combines Lempel-Ziv compression techniques with the Burrows-Wheeler transform but fails to index our data (see Section 3). Bloom filter trie (BFT) [17] method allows to index genome collections with a *k*-mer-based approach. *k*-mers are stored in a color-oriented graph, where each represents a set of potential *k*-mers. Furthermore, to each *k*-mer is associated a binary vector encoding its colors. These colors represent the genomes from which the *k*-mer is issued. AllSome sequence bloom trees (SBT) is an orthogonal approach to the BFT. SBT complexity scales up with the number of data sets [18]. A recent review on *k*-mer based approaches to store large collections of genomes can be found in [?].

### Our contribution

This paper provides a theoretical and practical contribution to the problem of finding a way to efficiently index large collections of similar genomes, assembled or not, without using information on variations from a multiple-genome alignment or a reference sequence. The data structure and the construction algorithm are described in Section 2. The time and space complexity are discussed in Section 2. Finally, the benchmark results of the current implementation of this algorithm, called RedOak, are provided in Section 3, followed by a discussion.

## 2 Results

The problem of indexing both assembled and unassembled genomes is equivalent to indexing a very large set of texts. This makes the problem related to the indexable dictionary problem, which consists of storing a set of words such that they can be efficiently retrieved [19]. A *k*-mer is a word of length *k* (a fragment of *k* consecutive nucleotides) of a read (sequence that came from high-throughput sequencing) or an assembled sequence (contig, scaffold, genome, or transcriptome). *k*-mers are words based on a simple alphabet Σ = {*A,C,G,T*}.

Before describing the way *k*-mers are indexed, we introduce some notation used in this paper. Given a set of *n* genomes 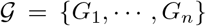, the *core k*-mers correspond to the subset, denoted 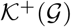, of the *k*-mers shared by all the genomes; the *shell k*-mers correspond to the subset, denoted 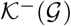, of the *k*-mers shared by at least one of the genomes but not by all. The set of all *k*-mers present in one or more genomes is denoted 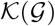 and is such that 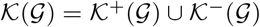. Given a prefix *pref* (of length *k*_1_ ≤ *k*), the subset of the *core k*-mers whose prefix is *pref* is denoted by 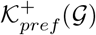, the subset of the *shell k*-mers whose prefix is *pref* is denoted by 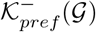, and the subset of all *k*-mers whose prefix is *pref* is denoted by 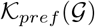. Given a *k*-mer *w*, we denote by 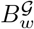 the Boolean array such that 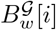 is *true* if and only if *w* occurs in the *i^th^* indexed genome (a.k.a., *G_i_*). In the remainder of the paper, the notation is shortened to 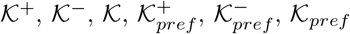, and *B_w_*.

There is a trivial bijection between the *k*-mers and their lexicographic rank. Because the alphabet is of size 4, only two bits (log_2_(4)) are required to represent each symbol. Let us assume that `A` is encoded by 00, `C` is encoded by 01, `G` is encoded by 10 and `T` is encoded by 11; any sequence of symbols of fixed length has a unique encoding scheme, which converts it into an unsigned integer that also represents its lexicographic rank among all the sequences of the same size.

To efficiently store and query the *k*-mers, each *k*-mer is split into two parts: its prefix of size *k*_1_ and its suffix of size *k*_2_, with *k*_1_ + *k*_2_ = *k*. Actually, the *k*-mers are clustered by their common prefix, and for each cluster, only the suffixes are stored. The choice of the value of *k*_1_ minimizing memory consumption is guided by both analytic considerations [20] and empirical estimation, as will be discussed in Section 3.

As described in Figure 1(a), the 4^*k*_1_^ clusters of *k*-mers are represented by an array of 4^*k*_1_^ objects (using their lexicographical order). The *i^th^* object corresponds to the set of *k*-mers whose prefix of length *k*_1_ is *i^th^* in the lexicographic order. Since the *k*-mers are grouped by common prefixes of length *k*_1_, there are 4^*k*_1_^ distinct clusters (array (1)). For each cluster, there are 4^*k*_2_^ possible suffixes (array (2)), which can be either absent from any of the indexed genomes (white cells) or present in some of the genomes *shell k*-mer (blue cells), present in one and only one genome *cloud k*-mer (orange cells) or present in all genomes *core k*-mer (green cells). When a *k*-mer is absent, all bits of its associated vector are set to 0 (array (5)). When a *k*-mer is present in all genomes, all bits of its associated vector are set to 1 (array (4)). In the last case, bits are set according to the presence/absence in each genome (array (3)).

**Figure 1:**
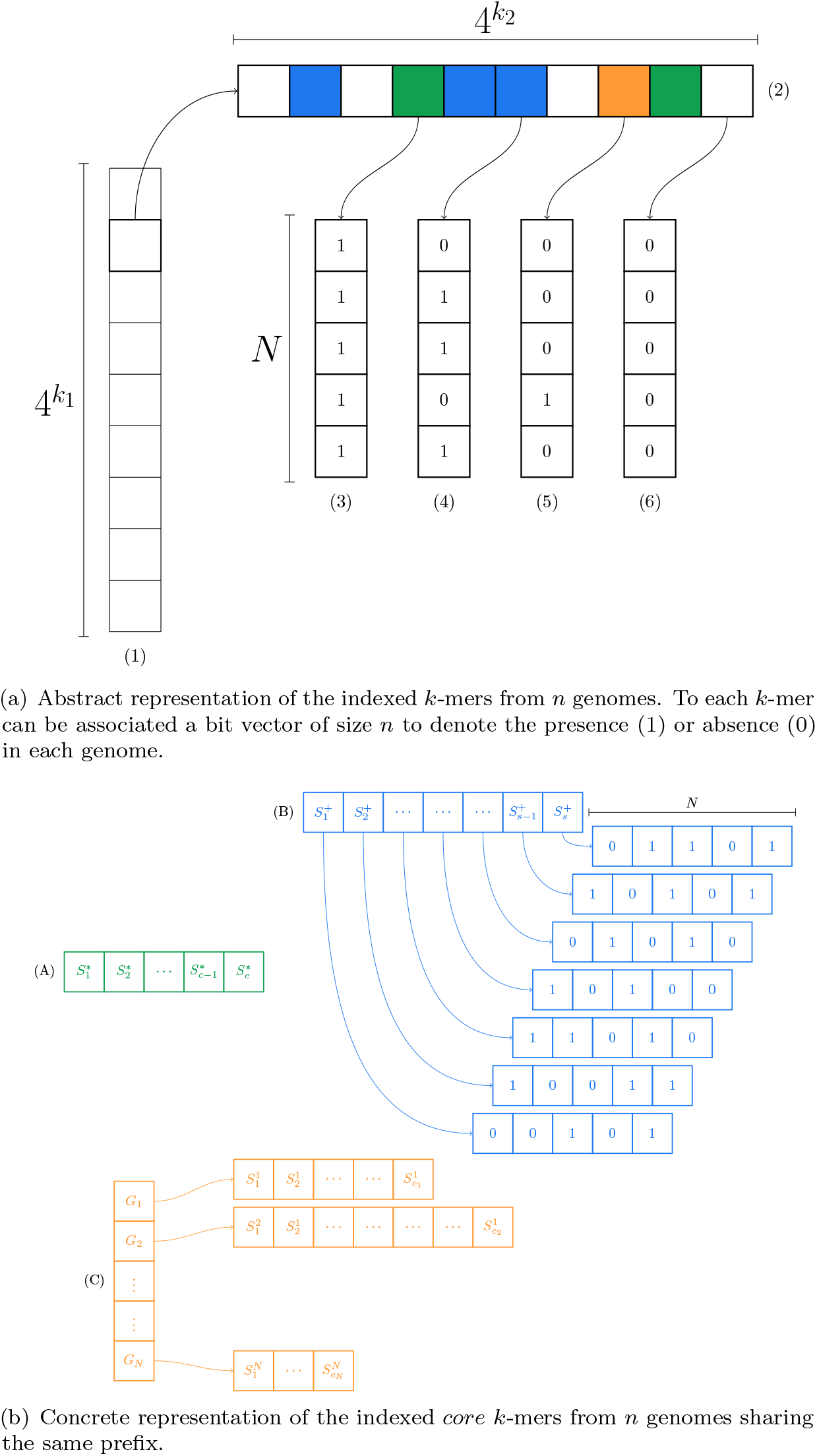
Representation of the data structure used to index the *k*-mers from *n* genomes. The (A) The green array represent *core k*-mers, (B) Blue cells the *shell k*-mers, and (C) in orange the *cloud k*-mers.

**Figure 2:**
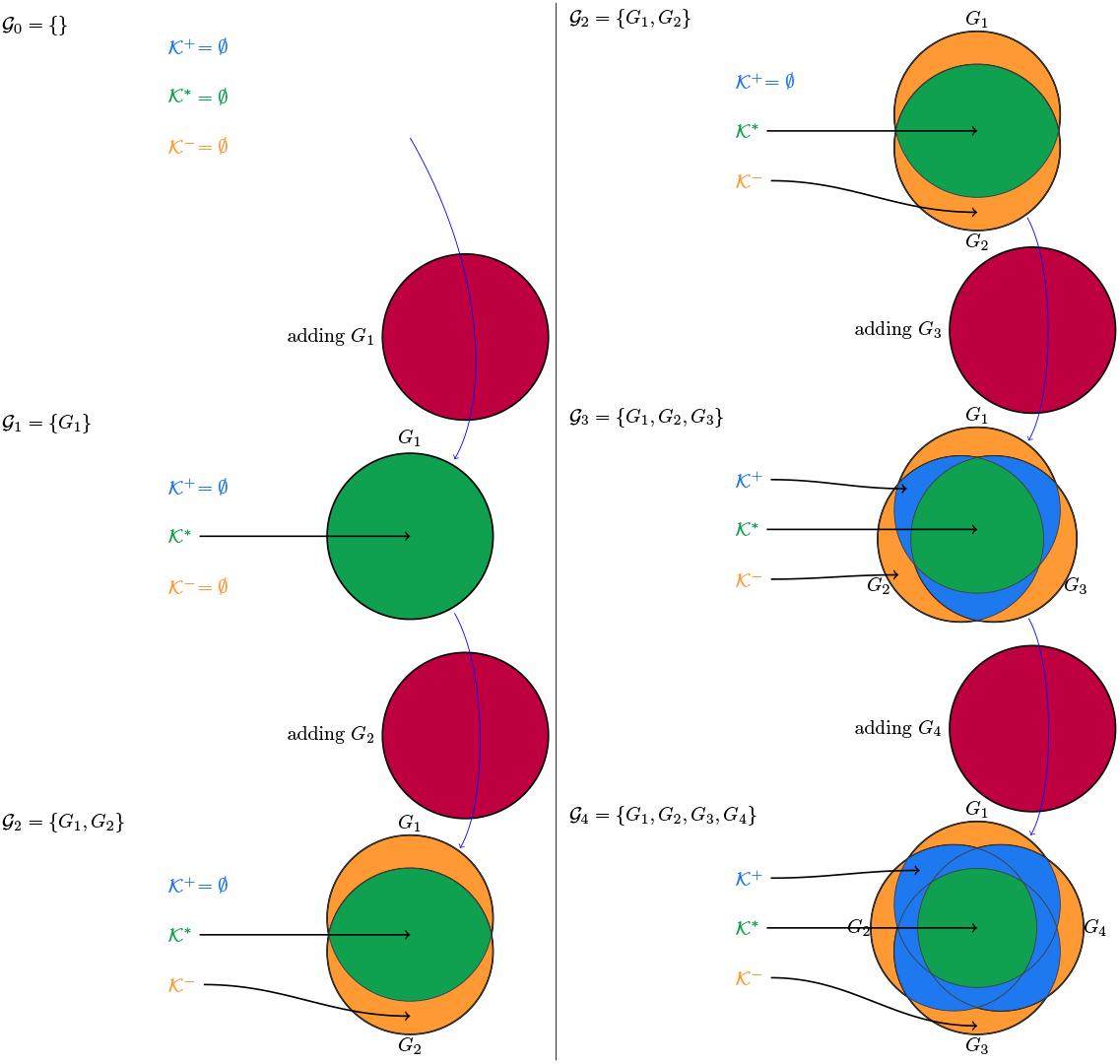
Illustration of the evolution of *k*-mers indexed by RedOak as the genomes are added.

On average, there are 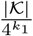 *k*-mers in each cluster. Even for small values of *k*_1_, this number is very low compared to the 4^*k*_2_^ possible suffixes. Thus, a bit-vector (even with a succinct data structure) cannot represent the array (2) (Figure 1(a)). Because *k*-mers not present in any genome (white cells in Figure 1(a)) are predominant and because they can be easily deduced from the other *k*-mers, they do not need to be explicitly stored. Moreover, a distinction is made between *core k*-mers (green cells in Figure 1(a)) and *shell k*-mers (orange cells in Figure 1(a)). Indeed, *core k*-mers are by definition present in all genomes and thus, it is not necessary to store information on which genome these *k*-mers are present.

The concrete representation of the data structure used to store the *k*-mers having the same prefix is shown in (Figure 1(b)). The *k*-mers absent from all genomes are obviously deduced from present *k*-mers and thus are not physically represented (and they all share the same 0-filled bit vector). The *k*-mers present in all genomes (*core k*-mers) are simply represented by a sorted vector where each suffix is encoded by its lexicographic rank (array (*A*)). These *k*-mers share the same 1-filled bit vector. The other *k*-mers (*shell k*-mers) are represented by an unsorted vector where each suffix is encoded by its lexicographic rank (array (*C*)). To each suffix is associated its presence/absence bit vector (array (3)). The order relationship between the suffixes is stored in a separate vector (array (*B*)). The *core k*-mers having the same prefix are stored in their lexicographic order (by construction) using 2 *k*_2_ bits, where *k*_2_ = *k* – *k*_1_ (array (*A*) of the Figure 1(b)). The *shell k*-mers are stored using 2*k* bits as well; however, their lexicographic order is not preserved (array (*C*) of the Figure 1(b)). Thus, this order relationship is maintained separately in another array, denoted 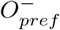 (array (*B*) of the Figure 1(b)). Moreover, for each represented *shell k*-mer *w*, a bit vector is associated with storing its presence/absence in the genomes (Figure 1(b), array (3), which represents *B_w_*).

In the RedOak implementation, both the *core* and *shell k*-mer suffixes are stored using 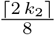 bytes each. The remaining unused bits are set to 0. This choice greatly improves the comparison time between *k*-mers suffixes. Moreover, because the presence/absence bit vectors are all of size *n* (the number of genomes), RedOak provides its own implementation for that structure, which removes the need to store the size of each vector. This implementation also emulates the 0-filled and 1-filled bit vector (arrays (4) and (5) of the Figure 1(a)).

The choice of this data structure was guided by the desire to allow genome addition without having to rebuild the whole structure from scratch. Indeed, indexing a new genome can be represented by some basic operations on sets as described in Listing 1. First, it is obvious that the only case where the set of *core k*-mers expands is when the first genome is added (line 11). The other updates of the *core k*-mers occur on lines 15 and 16 and only lead to the removal of some *k*-mers from this set.

Now, let us suppose that the set of *k*-mers of the new genome is lexicographic ordered (line 9). Then, the *core k*-mers are initially represented as a sorted (in lexicographical order) array, and it is easy to intersect these sorted *core k*-mers with the sorted *k*-mers from the new genome. During this step, there is no difficulty in producing, on the fly, both the subset of *k*-mers moving from the *core* to the *shell* (required at line 22) and the subset of *k*-mers from *g* that were not found in the *core k*-mers (required at line 24). Merging the elements coming from the *core k*-mers with the *shell k*-mers is equivalent to a concatenation of the two vectors (because no *k*-mer can be both *core* and *shell*); moreover, for each type, their associated presence/absence vector is 1-filled, except for the newly indexed genome. Merging the *k*-mers coming from the new genome requires that one first check if each “new” *k*-mer has already been indexed in the *shell*. In such case, the associated bit vector must be updated with the new indexed genome; otherwise, the new *k*-mer must be appended at the end of the *shell k*-mers with an associated 0-filled bit vector, except for the newly added genome. It does not matter which set of *k*-mers is appended; in both scenarios, the appended *k*-mers are sorted. Because the order relationship is stored for the old *shell k*-mers, it is easy to update the order relationship associated with the new *shell k*-mers by applying a trivial ordered set merging algorithm. This extra payload in memory enables faster processing than directly merging the *k*-mer suffixes and their bit vectors. Indeed, this auxiliary vector (array (*B*) of Figure 1(b)) uses 16-bit words (instead of 32 or 64 bits for pointers) to store the indices of the suffixes stored in 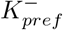 and the merging of the two orders.

Because the data structure partitions the set of indexed *k*-mers according to their common prefix of size *k*_1_, it is easy to parallelize the algorithm presented in Listing 1. Thanks to the Open-MPI specification [21], each instance of the RedOak program only processes a portion of the *k*-mers. This allows us to run RedOak on a cluster, on a multi-core architecture or on a combination of them. This feature has two major advantages: the required memory is split across the running instances, allowing scaling of the method to a very large collection of genomes, and the wall-clock time is drastically reduced (see Section 3).

Finally, the algorithm requires a strategy to output (in lexicographic order) all *k*-mers of each genome. The RedOak implementation is based on the `libGkArrays-MPI` (in prep.) library, which provides this feature.

The data structure presented in this section also has an interesting application: it enables easy and efficient queries. Querying for some sequence *s* consists of reporting, for all its *k*-mers, in which genome those *k*-mers appear. From that report, one can compute the number of *k*-mers of the query sequence that belong to each genome or the number of bases covered by the *k*-mers of the genomes (see Section 3). To query the data structure for a *k*-mer, the algorithm selects the *k*-mer prefix *pref* and then looks up (by dichotomy) its suffix in 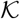 (specifically, in 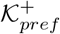 or 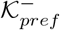). The time complexity is discussed in the next section.

In this part, we present the time and space complexity of the algorithm, using the notations below:

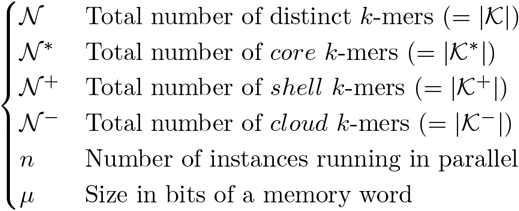

### Theorem 1.

*The space needed for indexing n genomes is equal to*

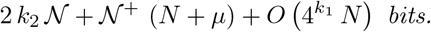

*If k*_1_ *is defined as* 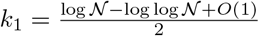, *then the memory space required by RedOak to index the k-mers of N genomes is increased by*

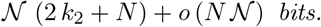

*Proof*. The structure associated with a *k*-mer prefix is identical to that described in Figure 1(b). Storing the suffixes of the *core* (resp., *shell* and resp., *core*) *k*-mers requires 2 bits per nucleotide, leading to 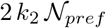 bits. In addition, a binary vector of size *N* and a memory word is associated with each suffix of *shell k*-mers, i.e. 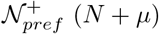 bits.

For a given prefix *pref*, the structure need 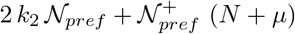 bits. To this must be added the data structures allowing to encapsulate information, thus representing *O*(*N*) octets.

Finally, a unique binary vector of size *N* is associated with *core k*-mers, just as a unique binary vector of size *N* is associated with *k*-mers absent from the structure as well as a unique binary vector of size *N* is associated with *cloud k*-mers of each genome *G_i_*(1 ≤ *i* ≤ *N*), totaling (*N* + 2) *N* + *O*(1) bits.

The memory space required by RedOak to index the *k*-mers of *N* genomes is therefore 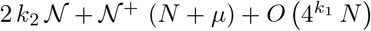 bits.

If 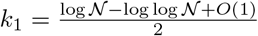, then 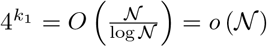.

It is also possible to notice that 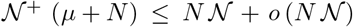. Thus, the memory space required by RedOak to index the *k*-mers of *N* genomes is therefore increased by 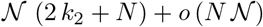 bits.

To this, we must add a space by node in *O*(1). However, it seems reasonable to consider that 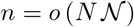.

Like the libGkArraysMPI library, for time performance reasons, we have chosen to use binary words of type `uint_fast8_t` for storing suffix information as well as binary arrays. Noting *μ*′ the number of bits of an integer of type `uint_fast8_t`, the space used for this storage is 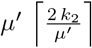 bits per suffix and 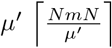 per binary array.

### Theorem 2.

*The time needed for indexing the* 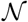 *distinct k-mers of n genomes is*

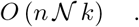

*Proof*. To study the data structure construction time, let us focus on the time required to add the set of *k*-mers sharing a common prefix *pref* (coming from a new genome *G*_*n*+1_) into an existing index of *n* genomes. Denote this set by *K_pref_* and its size as *M_pref_*. Assume that this set is already in lexicographical order. Computing the intersection between 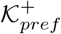 (sorted by construction) and *K_pref_* requires 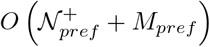 suffix comparisons. Suffix comparison requires 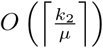 operations. Each time a suffix from *K_pref_* is found in 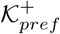, it is removed (the next suffixes from to be retained will be shifted back in the array by as many removed suffixes). Each time a suffix from 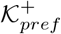 is not found, it is moved into a new temporary array (the next suffixes from 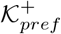 will be shifted back in the current array by as many removed suffixes as well). Shifting a suffix requires 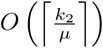 (operations. Thus, for this step, the overall time complexity is 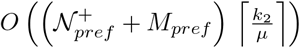. For speed optimization, the RedOak implementation pre-allocates an array for the suffixes to move from 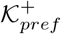 to 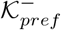 of length 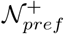, which is on average 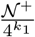.

Let 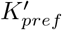 be the *k*-mer suffixes not found in 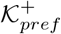, and let 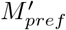 be the number of such *k*-mers. Computing the union of 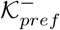 with 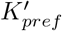 is not significantly more difficult. First, all bit vectors must be extended, which could be costly; however, the capacity of the bit vectors can be allocated beforehand, leading to a constant operation for this extension. The RedOak implementation computes the total number of genomes before indexing them. Therefore, by default, any *k*-mers in 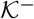 should be absent in the new genome being currently added. It follows that each suffix *suff* from 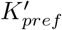 is searched in 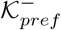. If it is found, then the last bit of *B_pref·suff_* is set to 1 and is removed from 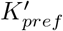. Since the order relationship of 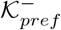 is retained separately in a specific array 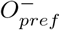, this step requires 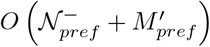 suffix comparisons. The remaining suffixes from 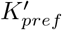 (say, 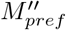) are then appended to the end of the array storing 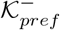. For each suffix, its bit vector is added. This step requires 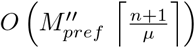 operations. Furthermore, the order array 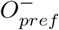 is extended to consider the newly added suffixes, and the reordering requires 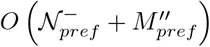 operations. Ultimately, since this operation is performed for every prefix, the overall time complexity of adding an ordered set of *M k*-mers to the current index is 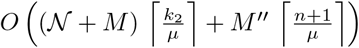.

To this complexity, the time for producing the ordered set of *k*-mers grouped by suffix should be added. RedOak uses the libGkArrays-MPI implementation, which, assuming that 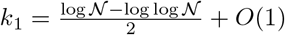, runs in *O* (*k M* log log *M*). It is obvious that the number of distinct *k*-mers is bounded by the size of the added genome. Assuming that both 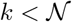, *M* log log 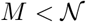, the total running time for adding *n* genomes of size *m* is bounded by 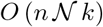.

### Theorem 3.

*Assuming that the number of genomes per indexed k-mer follows a Poisson distribution of parameter* λ (*where* λ *is the average number of genome sharing a k-mer), the size of* 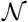 *is*

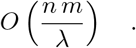

*Proof*. Since the run time clearly depends on the number of indexed *k*-mers, let us use a simple model to approximate the time complexity. Suppose that each genome has *m* distinct *k*-mers and that each *k*-mer has a fixed probability *p_i_* to be shared exactly by *i* genomes out of *n*. The total number of indexed *k*-mers is then

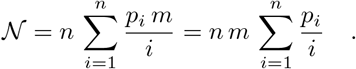

In the worst case, each *k*-mer is specific to each genome (*p*_1_ = 1), which leads to 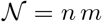. In contrast, the best case occurs when all *k*-mers are *core*. In such a situation, 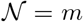. If all 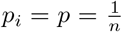, since 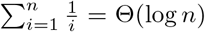, then 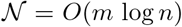. Now refine the model and denote by λ the average number of genomes sharing the indexed *k*-mers (1 ≤ λ ≤ *n*). The probabilities *p_i_* then follows a Poisson law of parameter 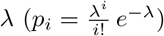. Thus,

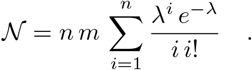

Let us recall 1/ that the exponential integral 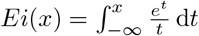 is such that

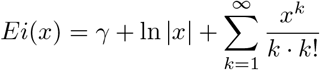

(where *γ* is the Euler-Mascheroni constant) and 2/ that the logarithmic integral 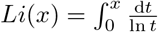 (for *x* ≠ 0) is such that

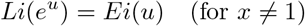

and *Li*(*x*) behaves asymptotically for *x* → ∞ to 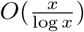.

Bounding 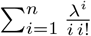 by 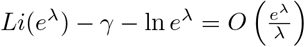 gives:

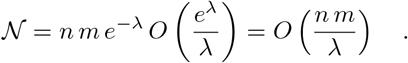

### Theorem 4.

*Given the index of* 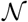 *k-mers from n genomes with* 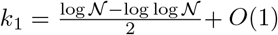, *querying the index for all k-mers from a sequence s requires* 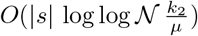 *operations*.

*Proof*. Extracting the first *k*-mer of the query *s* (and computing its prefix *pref* of size *k*_1_) requires *O*(*k*) operations. Extracting the other *k*-mers (and computing their prefix) can be performed in *O*(1) operations for each. Thus extracting all the *k*-mers of the query sequence requires *O*(|*s*|). As already stated above, for each *k*-mer, there are on average 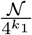 suffixes associated with its prefix; thus, performing a dichotomic lookup requires, on average, 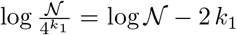 comparisons between suffixes. By choosing an appropriate value of 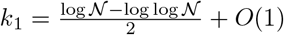, the number of lookups become 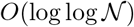.

### Theoretical predictions

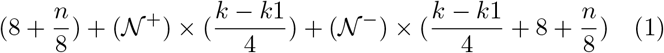

### Simplified theoretical cost

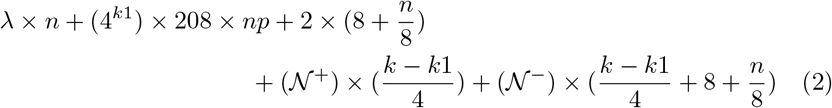

### Estimated cost

For 67 genomes with 40 instances, *k* = 27, k1 = 12 with 10% of core *k*-mers, and 90% of dispensable *k*-mers:

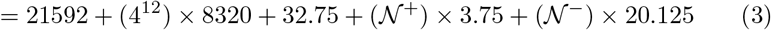

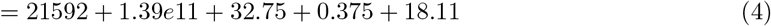

### Estimated cost per nucleotide

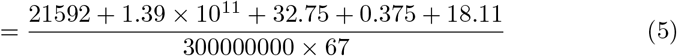

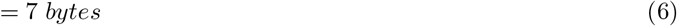

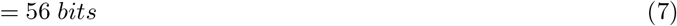

In (4), λ is the Index class size, (152) bytes, plus the Genome class size, (320) bytes, times the number of genomes. Theoretically (according to (7)), we only need 56 bits; however, in practice, we use 35 bits per nucleotide for 67 assembled genomes indexed.

## 3 Discussion

### Implementation

RedOak is implemented in C/C++ and its construction relies on parallelized data processing. A preliminary step, before indexing genomes, is performing an analysis of the composition in *k*-mers of the different genomes. During this step, *k*-mer counting tools could be involved and their performance is crucial in the whole process [22]. We looked for a library allowing us to handle a large collection of genomes or reads, zipped or not, working in RAM memory, and providing a sorted output. Indeed, RedOak uses libGkArrays-MPI^1^ which is based on the Gk Arrays library [23]. The Gk array library and libGkArrays-MPI are available under CeCILL licence (GPL compliant).

The libGkArrays-MPI library is highly parallelized with both Open MPI and OpenMP.

To manipulate *k*-mers, the closest method is Jellyfish [24]. This approach is not based on disk but uses memory and allows the addition of genomes to an existing index. However, we did not use it because in the output, *k*-mers are in “fairly pseudo-random” order and “no guarantee is made about the actual randomness of this order”^2^.

### Value of *k* and *k*_1_

In most of the *k*-mer based studies, the *k*-mer size varies between 25 (with reference genome) and 40 (without reference genome). The value of this parameter can be statistically estimated as stated in [25].

The *k*_1_ prefix length in our experiments has been defined on the basis of analytic considerations presented in [20] but can be arbitrarily fixed to some value between 10 and 16, which respectively leads to an initial memory allocation from 8MiB to 32GiB, equally split across the running instances of RedOak. Setting a higher value is not necessary; otherwise, it may allocate unused memory.

### Benchmark

The experiments were performed on a SGE computer cluster running Debian. The cluster (SGE 8.1.8) has two queues. The “normal” queue has 23 nodes, having 196 GiB of RAM and 48 cores^3^ each. This queue represents 4.4 TiB of memory and 1104 cores. The “bigmem” queue possesses 1 node, having 2 TiB of RAM and 96 cores^4^. The benchmark was performed on both “bigmem” and “normal” queues. Our dataset of 67 uncompressed rice genomes is equal to 25 Gib. which represent 26194967769 nucleotides.

### Comparison of RedOak, Jellyfish and BFT for the index build step

We compared RedOak to two other methods, namely Jellyfish [24] and Bloom Filter Trie (BFT) [17]. The comparison was performed on the 67 *de novo* assembled rice genomes from Zhao et al. [26] by comparing the time used for the index build phase and the maximum memory consumption. The size of the data set was successively set to 10, 20, 30, 40, 50, 60 and 67 genomes out of the original data set.

Jellyfish builds an index for each genome, and then these indexes were merged to produce a matrix where the counts for each *k*-mer in each genome are stored (small modification of the merge tool implementation of Jellyfish). For JellyFish, we also created a program that simulates a parallelization of jobs.

BFT needs ASCII dumps to build its index. These dumps were produced using Jellyfish. For Jellyfish and BFT the reported values are the total time taken for both the counting and merging steps. For all experiments, the *k*-mer size was set to *k* = 27, since BFT requires *k* to be a multiple of 9. For RedOak, the prefix length was set to 12 (default setting), which gives a table of prefixes of very reasonable total size *i.e*., 4^12^ = 128 MiB. Each prefix index of each running instance represents 3.2 MiB *i.e*., 32 MiB by node. This drastically reduces the risk of saturation during the experiments.

For each subset, we used RedOak in parallel on 10 “normal” nodes of the cluster and on each node we reserved 4 cores. For each subset, we also used JellyFish (jellyfish count -m 27 -s 500 M -t 10) on 40 genomes in parallel using 40 nodes. BFT does not allow merging the indexes created and does not propose parallelization. Therefore, we ran each instance of BFT in parallel using one “bigmem” node for each subset.

The results are summarized in Table 1 and in Figure 3. BFT was not able to index datasets in the runs with 40 or more genomes. Overall, RedOak showed better performance compared to JellyFish. RedOak used 2GiB per instance, and because it is parallelized on 40 instances, it used 80GiB for all the subsets and for the 67 assembled genomes. The index construction time in second was constant at approximately 1467.8 sec per ten genomes and took a total time of 8092.8 sec for the 67 genomes.

**Table 1:**
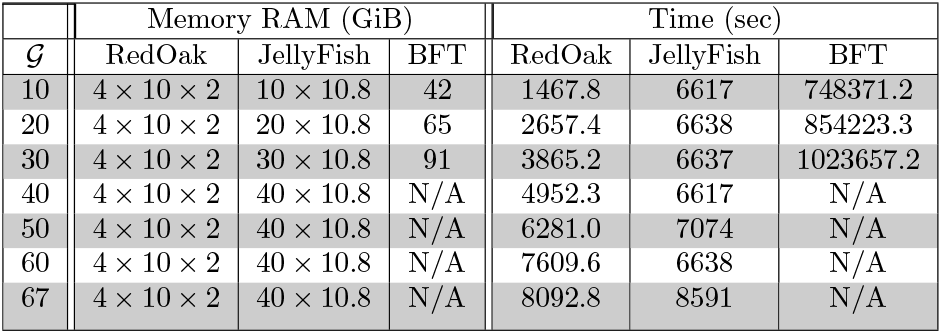
Performance comparison between RedOak, Jellyfish and BFT for the index build step. The size of the input was successively set to 10, 20, 30, 40, 50, 60 and 67 assembled genomes. RAM usage is in GiB. Times shown are wall-clock run times in sec.

**Figure 3:**
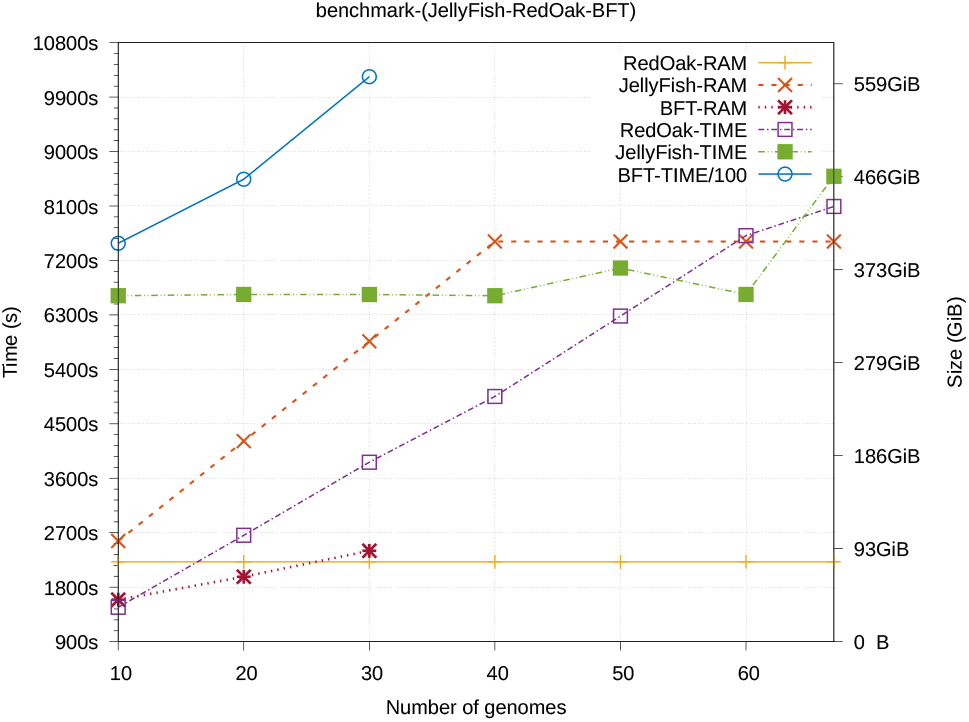
Performance comparison between RedOak, Jellyfish and BFT for the index build step. The size of the data set was successively set to 10, 20, 30, 40, 50, 60 and 67 genomes out of the original data set (x-axis). A dot represents the wall-clock run time (y-axis) or the RAM usage (y2-axis) required to build the index. The colors represent the softwares used: RedOak, Jellyfish or BFT. For BFT, we divided the construction time by 100 to fit our figure.

### Query performance

We also assessed the performance of RedOak for querying with sequences of different lengths the index of the 67 assembled rice genomes. We compared RedOak and JellyFish using random query sequences of length varying from the size of 10 times the size of *k* to 1000 times the size of *k*. The results are presented in Table 2, showing the maximum RAM usage and wall-clock run time required to match the 67 assembled rice genomes with a randomly created sequence.

**Table 2:** Comparison of performance between RedOak and JellyFish for querying with simulated sequences of different length (from 10k to 1000k) an index of 67 assembled genomes. Maximum RAM usage are in GiB. Times shown are wall-clock run times in sec.

| Query length | Memory RAM (GiB) and Query Time (s) |  |  |  |  |
| --- | --- | --- | --- | --- | --- |
|  | RedOak-RAM | RedOak-Query | JellyFish-RAM | JellyFish-Query | JellyFish-Query-Total |
| 270 | 7 | 1.179 | 16 | 6360 | 257926 |
| 540 | 7 | 3.325 | 16 | 7919 | 301113 |
| 810 | 7 | 2.703 | 16 | 11586 | 374489 |
| 1080 | 7 | 6.847 | 16 | 12996 | 368945 |
| 1350 | 7 | 3.839 | 16 | 12280 | 351517 |
| 2700 | 7 | 9.006 | 16 | 12880 | 391060 |
| 5400 | 7 | 19.634 | 16 | 13397 | 337673 |
| 8100 | 7 | 24.976 | 16 | 11701 | 349188 |
| 10800 | 7 | 34.939 | 16 | 10668 | 262252 |
| 13500 | 7 | 43.997 | 16 | 9779 | 263889 |
| 27000 | 7 | 60.389 | 16 | 9569 | 295048 |

To evaluate the query time of JellyFish, we had to request each file individually. The Table 2 shows, for JellyFish, the max RAM memory (including the writing time of all *k*-mers), the time of the longest query, and the total time (sum of all times which gives us the average time: 4403.7 sec per file). The results showed that RedOak has better performance than JellyFish for querying this dataset.

### Example of PAV analysis

Analysis of presence–absence variation (PAV) of genes among different genomes is a classical output of pan-genomic approaches [1], [27], [26]. RedOak has a nucleotide sequence query function (including reverse complements) that can be used to quickly analyze the PAV of a specific gene among a large collection of genomes. Indeed, we can query, using all *k*-mers contained in a given gene sequence, the index of genomes. For each genome, if the *k*-mer is present in any direction we increment the score by 1. If the *k*-mer is absent but the preceding *k*-mer (overlapping on the first *k* – 1 nucleotides) is present, we note that there is an overlap, but RedOak does not increase the score. If the score divided by the size of the query sequence is greater than some given threshold, then we admit that the query is present in the genome.

As an example, we indexed the 67 rice genomes from Zhao et al. [26] with RedOak using *k* = 30, and we accessed the PAV of all the genes from *Nipponbare* and one gene from *A. Thaliana* using a threshold of 0.9. the gene Pstol, which controls phosphorus-deficiency tolerance [28]. For a specific genome (GP104), we were able to detect the gene presence of the gene Pstol, whereas this presence has not been found in Zhao et al. [26].

We need to keep in mind that this score under-estimates the percentage of identity. Indeed, let us suppose that the query sequence (of length *ℓ*) can be aligned with some indexed genome with only one mismatch, then all the *k*-mers (of the query) overlapping this mismatch may not be indexed for this genome. This implies that only one mismatch can reduce the final score by 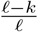, whereas the percentage of identity is 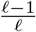. Said differently, in this experiment, a query having a score ≥ 0.9 can potentially be aligned with a percentage of identity greater than 97%

### Indexing a collection of unassembled genomes

We accessed CHICO [16] on a set of FASTQ files extracted for only 10 genomes from the 3000 rice genomes project [29] and ran out of memory. Using the reads from zipped FASTQ files, RedOak was able to index a subset of 110 randomly chosen, unassembled genomes from the 3000 rice genomes project. It ran 140 parallelized instances on 14 nodes, each using 10 cores. It used a total of 47254459 sec and 683.337 GiB of RAM memory. Per instance, it used 337531.85 sec (4 days) and 4.881 GiB.

## 4 Conclusion

We have designed and developed a data structure dedicated to the indexation of a large number of genomes, assembled or not. The parallelization of the data structure construction allows, through the use of networking resources, to efficiently index and query those genomes.

Several perspectives can be considered. Through intensive tests and scalability proofs, we aim to guarantee the robustness of our approach and extend the usability of our tool, for instance, by proposing a graphical interface.

We also can explore methods inspired by Bloom Filter Trie [17], using a probabilistic approach. At each vector level, there are possibilities to introduce such a model. A theoretical study must be performed to estimate the possible gains and losses of such a model.

We also have in mind other uses of the matrices of *k*-mers to extend the applications of our data structure. For instance, beyond genomes comparison, we could also consider using them for analyses of variant detection, genome assembly improvement, and phylogeny.

## Availability and requirements

Project name: RedOak

Project home page: https://gite.lirmm.fr/doccy/RedOak

Operating system(s): Platform independent

Programming language: C++

Other requirements: See webpage (Readme.md)

License: Creative Commons

Any restrictions to use by non-academics: none

## Declarations

### Ethics approval and consent to participate

Not applicable

### Consent to publish

Not applicable

### Availability of data and materials

The source code of RedOak is freely available at the repository https://gite.lirmm.fr/doccy/RedOak.

### Author’s contributions

CA, GD, GS and AM implemented and tested the methods. CA, AC, AM and MR designed the methods, the experiments, and the analysis. CA, AC, AM and MR wrote, edited, and revised the paper.

## Acknowledgements

Our thanks go to the members of the GenomeHarvest project members, and especially to Anne Dievart and collaborators for their implications in discussion and having provided their data.

## Listing 1: High level algorithm to incrementally update the index

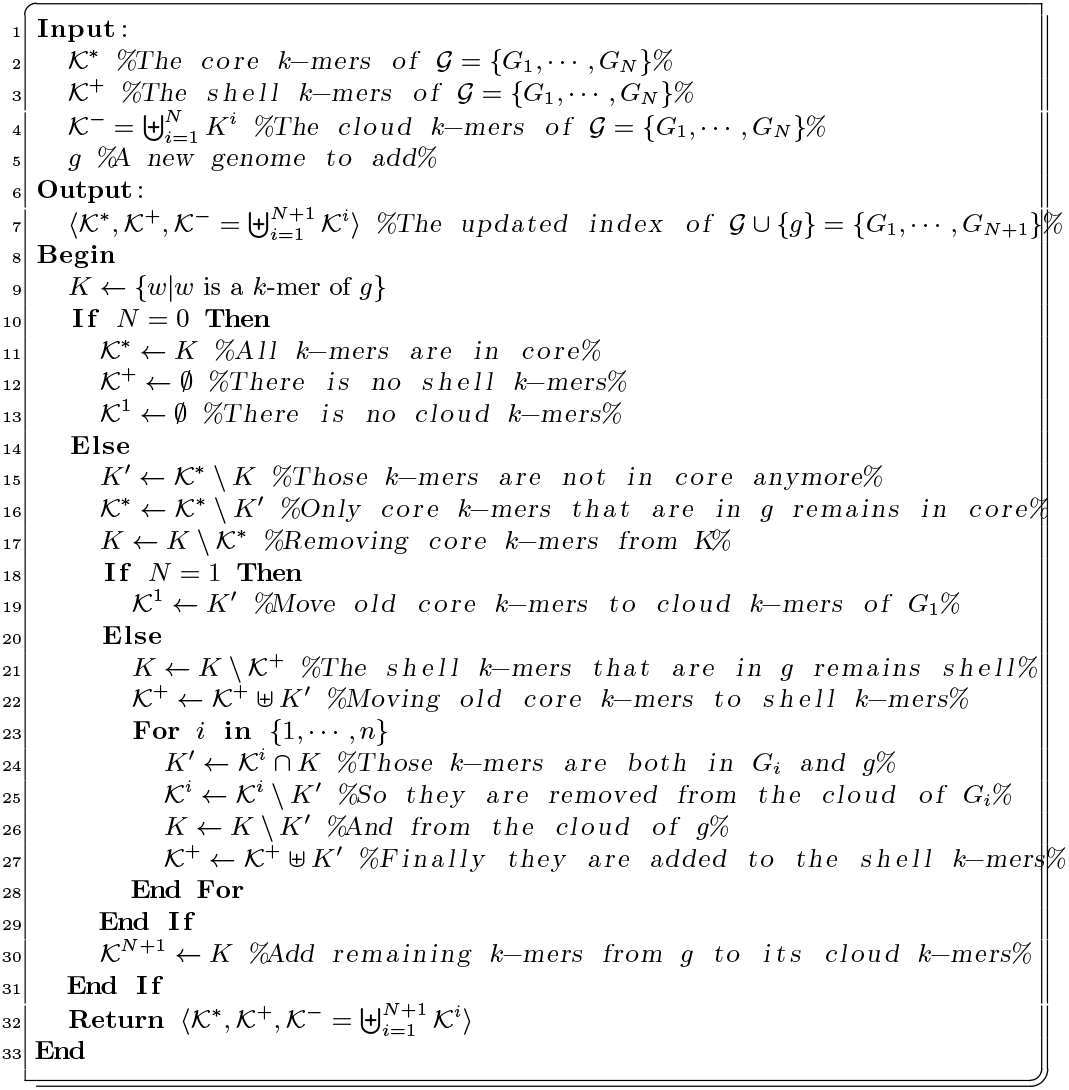

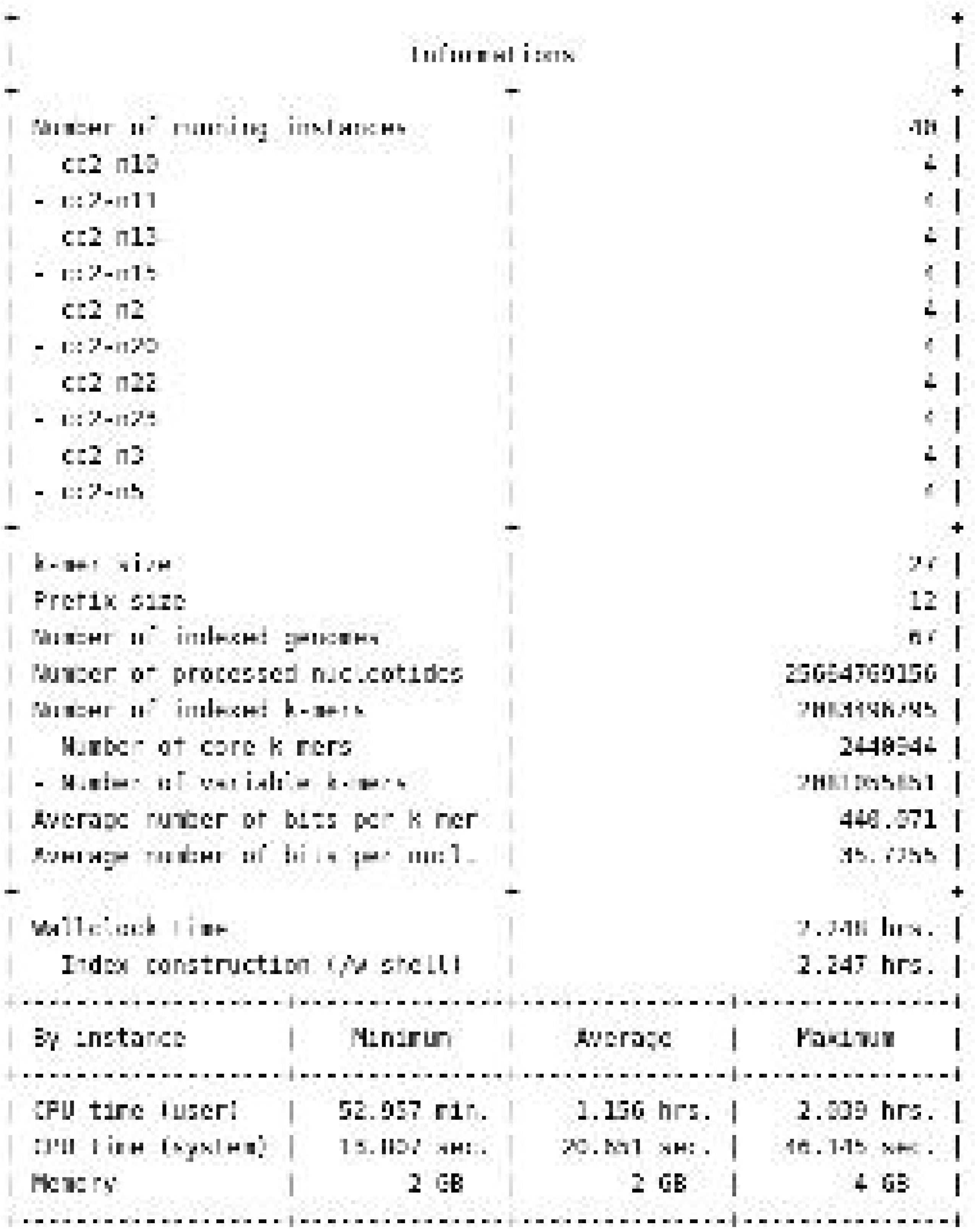

## Footnotes

1 Private communication, Mancheron *et al*..

2 Documentation of JellyFish.

3 Intel^®^ Xeon^®^ CPU E5-2680 v3 processor clocked at 2.50GHz.

4 Intel^®^ Xeon^®^ CPU E7-4830 v3 processor clocked at 2.10GHz.

